# Development and clinical validation of molecular subgrouping in medulloblastoma by targeted methylation sequencing

**DOI:** 10.1101/2023.12.08.570900

**Authors:** Shreya Srivastava, Kamlesh Bhaisora, Naveen Kumar Polavarapu, Lily Pal, Shalini Singh, Neha Rai, Raghavendra Lingaiah

## Abstract

**Background:** The WHO classification of CNS tumors confers promising prognostic value to the molecular classification of medulloblastoma (MB). Next-generation sequencing (NGS) has been the primary method employed for molecular classification through transcriptomic, genomic, or methylation profiling. However, due to cost and infrastructural needs, particularly in developing countries, we propose a relatively simple, rapid, and economical Sanger sequencing-based targeted methylation sequencing method for MB classification and prognostication.

**Methods:** Eleven epigenetic targets were amplified using optimized primers and bisulfite-converted DNA for Sanger sequencing. Chromas software was used for low-quality data trimming and NCBI’s Needleman Wunsch alignment tool was used for sequence alignment to reference. The developed method was applied to tissues from twelve cases of medulloblastoma.

**Results:** Successful interpretation of methylation status in ten out of eleven targets was achieved which was sufficient for classification according to the latest WHO classification of Medulloblastoma tumors. Twelve medulloblastoma cases were classified into WNT (n=2), Group 3 (n=5), and Group 4 (n=5).

**Conclusion:** The developed Sanger sequencing method is a cost-effective, in-house solution that can be used for molecular subgrouping of medulloblastoma. It offers an alternative to NGS, can be done on a case-to-case basis, and does not require high-end infrastructure, sample pooling, or extensive bioinformatics knowledge.

**Impact statement:** Molecular classification is imperative for determining the prognosis of medulloblastoma and is recommended by WHO. However, NGS proves to be an expensive option in developing countries. This study has come up with an affordable targeted methylation Sanger sequencing method requiring minimal bioinformatic skills, by utilizing epigenetic targets, for prognostication and risk stratification in medulloblastoma patients. The molecular subgroups of all recruited cases were successfully determined according to WHO classification which is crucial information that, when combined with clinical findings, will enable the clinicians to determine effective treatment strategies.

## 1. Introduction

Paediatric brain tumors are the second most common type of cancer in children, after leukemia (1). Tumors of the central nervous system (CNS) make up about 2% of all cancers, with Medulloblastoma (MB) being the most common, accounting for 12-25% of all CNS tumors (2–6). It is a malignant tumor of the posterior fossa affecting children between 1-9 years of age, with surgery, radiation, and chemotherapy as the primary treatment strategies. The World Health Organization (WHO) classifies tumors of the CNS into distinct subtypes based on histological and molecular features. MB is classified histologically into classic, desmoplastic/nodular, MB with extensive nodularity, and large cell anaplastic subtypes, while molecularly it is subtyped into WNT, SHH, and Non-WNT/Non-SHH groups, which are further subdivided into Group 3 and Group 4 tumors (7). The recently updated WHO classification (2021) emphasizes the molecular classification of MB as it provides clinical and prognostic significance beyond that provided by histopathological examination where molecularly defined Group 3 has a worse prognosis while WNT shows the best prognosis (8). WHO also recommends an integrated diagnosis based on histology, grading, and molecular/genetic information (9).

However, in most clinical settings, MB diagnosis, risk stratification, and treatment solely rely on histological, radiological, and clinical findings. This results in a vague categorization of patients into risk groups, making it difficult to prognosticate and manage patients. Molecular subtyping can help in MB prognostication, risk stratification, and deciding treatment modalities.

Earlier studies utilized whole genome sequencing and gene expression profiling for molecular classification, but shared features among subgroups failed to clearly define subtypes. Currently, epigenetic markers related to DNA methylation are widely used to classify MB based on reports of mutations in epigenetic regulators (10,11).

Although there are various platforms for methylation profiling of medulloblastoma (MB) for molecular classification, their feasibility, and applicability remain questionable across all clinical settings. In this study, we developed an in-house method for targeted methylation sequencing based on Sanger sequencing, using epigenetic targets previously reported by Gomez et al., and evaluated its performance on formalin-fixed paraffin-embedded (FFPE) and fresh tissues.

## 2. Method

### 2.1 Patients and sample collection

This study utilized tissues from patients with confirmed Medulloblastoma diagnosis and was ethically approved by the Institutional Ethics Committee (IEC code-2020-316-IMP-EXP-33). Two FFPE tissue blocks (12-16 months old) were obtained from the Department of Pathology and twelve fresh tumor tissues were collected prospectively at the time of surgery from the Department of Neurosurgery. Fresh tissues were collected in 500μl 1X Tris EDTA (TE) buffer and stored at –80L until DNA extraction. Figure 1 outlines the methodology.

**Figure 1:**
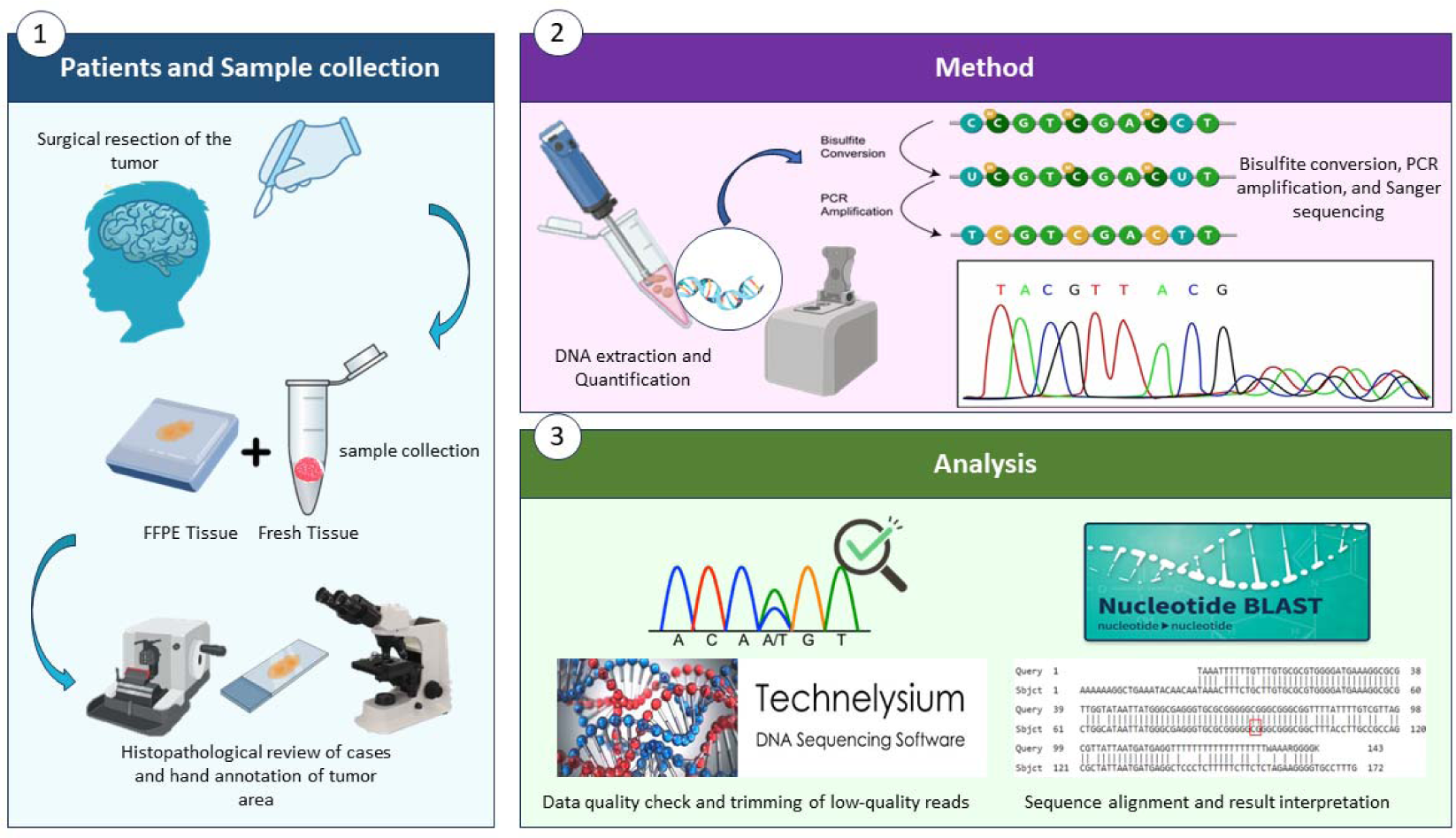
Sequential study design.

### 2.2 Primer designing

Eleven primers were designed using UCSC Genome Browser and MethPrimer Plus software from the Li Lab (Urogene.org) against CPG islands previously reported by Gomez et al. (18) in the following genes (Illumina array IDs in brackets): LHX6 (cg25542041), CHTF18 (cg10333416), USP40 (cg12925355), AKAP6 (cg18849583), KIAA1549 (cg01268345), VPS37B (cg13548946), RPTOR (cg09929238 and cg08129331), RIMS2 (cg12565585), and two Illumina Array IDs – cg02227036 and cg05679609 with hg19 as reference. The amplicon length ranged from 200-250 bp, except for USP40 and RPTOR with amplicon lengths of 167 and 397 bp, respectively.

### 2.3 Tissue processing and DNA extraction

PCR optimization was done on DNA extracted from a healthy donor’s blood using a DNeasy kit (Qiagen, Germany). For fresh tissue, 20-25mg tissue cut from the tumor area was homogenized using a hand-held tissue homogenizer (MT hand-held homogenizer) and incubated at 56LJ until complete lysis (2-3 hours) before DNA extraction by PureLink Genomic DNA mini kit (Invitrogen™). For FFPE tissue processing and DNA extraction, the tumor area in H&E slides of selected cases was assessed and hand-annotated by a Neuropathologist. DNA extraction was performed as per kit protocols of the QiAamp FFPE DNA kit (Kit1) and Promega Reliaprep FFPE gDNA Miniprep system (Kit 2). 5×10μm thick sections were sliced from the FFPE tissue block and collected on egg albumin-coated slides. Contrary to mineral oil treatment for deparaffinization in Kit 2, Xylene as a dewaxing agent was used in the Kit 1 protocol. Using a decreasing gradient of ethanol tissue was rehydrated and an area parallel to the marked area on H&E slides was scraped off and collected in a sterile 1.5ml microcentrifuge tube to which 180µl tissue lysis buffer and 20µl Proteinase K were added and incubated at 56LJ until complete digestion.

### 2.5 Quantitative and qualitative evaluation of DNA extracted

The DNA extracted from blood, FFPE, and fresh medulloblastoma tissue was quantified using the Nanodrop spectrophotometer. For the qualitative evaluation of DNA, 2µl of DNA from xylene-treated and mineral oil-treated FFPE tissues and fresh tissue was loaded on 1.5% agarose gel stained with ethidium bromide, to check for DNA integrity.

### 2.6 Bisulfite conversion

As the primers designed are targeted against methylated DNA regions, the DNA extracted from FFPE/Fresh tissue as well as blood was bisulfite converted using a commercially available kit-the Zymo EZ Methylation Gold and Zymo EZ Methylation Lightening kit from Zymo Research. A controlled DNA input of 200-500ng in 20μl at the bisulfite conversion step was ensured to achieve maximum conversion efficiency and avoid incomplete conversion. The bisulfite-converted DNA was quantified again as ssRNA using Nanodrop Spectrophotometer to check for DNA recovery post-conversion.

### 2.7 PCR optimization

Primers for the aforementioned epigenetic targets were optimized for their annealing temperature with an input of bisulfite-converted control blood DNA in gradient PCR using four temperatures 3LJ to 5LJ lower than their melting temperature. Optimized PCR conditions were used to amplify epigenetic markers from bisulfite-converted DNA from blood, FFPE, and fresh tissues (20-30ng). Successful amplification was verified by running the PCR products on 3% ethidium bromide-stained agarose gel.

### 2.8 Sanger sequencing

An enzymatic cleanup using ExoSap (Applied BiosystemsTM) was done on PCR products before sequencing to remove unused dNTPs and primer dimers. Using the BigDye terminator cycle sequencing kit (Applied Biosystems^TM^), reactions were set up for each sample using forward primer for eleven distinct targets followed by post-cycle sequencing cleanup by BigDye X terminator purification kit (Applied Biosystems^TM^).

Sequencing was carried out on a 3500×l Genetic Analyzer from Applied Biosystems (ABI).

### 2.9 Data quality check

The quality of. ab1 chromatogram files from Data Collection 3500 software (ABI) was assessed in Seq A6 software (ABI). The KB™ Basecaller provided a quality value (QV) prediction per base that represented the probability of base call error and mixed base calls, with QV>20 denoting a 1% rate of error. The sequencing reads were checked for their QV values and signal intensity of approx. or >1000 relative fluorescence units (RFU values). Noisy data was excluded by investigating the average signal-to-noise ratio for four bases per sample. The read length in sequencing data was determined by comparing the bases called by the sequencer with the amplicon length of the respective targets. Sequencing reads with low signal intensity, low signal-to-noise ratio, truncated sequencing reads, or failed sequencing was sequenced again.

### 2.10 Sequencing Alignment and Analysis

Methylation analysis of eleven epigenetic markers was done by aligning sequencing data with the non-bisulfite-treated reference gene sequence for each target. ab1 chromatogram files were imported into Chromas software (Technelysium) for trimming low-quality bases and exported as FASTA files. Aligned to the original unmethylated sequence of target regions using NCBI’s Global Alignment Tool – Needleman-Wunsch alignment, methylation status was inspected by estimating the presence of Cytosine or Adenine at the target site in reference to the nucleotide present at the same site in the original unmethylated reference sequence.

### 2.11 Data interpretation

For Data Interpretation, epigenetic classifiers from a study by Gomez Et al. consisting of Classifier 1(^Epi^WNT-SHH) comprising six epigenetic targets, and Classifier 2 (^Epi^G3-G4) with five epigenetic targets were used as a reference for the analysis of methylation status. While, Classifier 1 was efficient in subgrouping medulloblastoma cases into WNT, SHH, or Non-WNT/Non-SHH groups, it was incapable of classifying Non-WNT/Non-SHH groups into Group 3 and Group 4 tumors. For this, cases with determined Non-WNT/Non-SHH groups were further tested for the five epigenetic targets belonging to classifier 2, for their subgrouping into either Group 3 or Group 4.

### 2.12 Correlation with histological and radiological findings

Pre-operative radiological and post-operative histopathological findings were collected for all recruited cases. Molecular pathologists remained blinded to clinical findings until the final analysis. Subgroups determined from a developed method were correlated with radiological features and histological type.

## 3. Results

### 3.1 DNA extraction and bisulfite conversion

A suitable quality and quantity of DNA was obtained from both blood and fresh tissues, which was markedly higher than FFPE tissues. In the case of FFPE tissues, a notable effect of the deparaffinization agent on DNA yield was observed. DNA extraction by kit 1 post xylene treatment resulted in a relatively low DNA yield perchance due to DNA degradation at the deparaffinization step in addition to tissue fixation with formaldehyde and A260/280 purity ratio > 1.8 as a consequence of RNA contamination.

Therefore, the use of mineral oil for deparaffinization was implemented in the protocols of Kit 2 to prevent degradation that significantly increased the DNA yield and ideal A260/280 ratios were achieved using RNAse. In addition, mineral oil helped restrain the high lipid content in the DNA extracted, a feature of brain tissues, that might interfere with the column-based DNA extraction methods resulting in low DNA yield or reduced amplification during PCR. Due to the solubility of lipids in mineral oil, a 6-7-fold increase in DNA was obtained compared to DNA extraction post-xylene treatment (See Supplemental Table 1).

The integrity of DNA obtained after deparaffinization by either xylene or mineral oil was checked on a 1.5% agarose gel. Smearing of bands was observed in both cases, possibly due to fragmentation caused by fixation and embedding of tissues (See Supplemental Figure 1). Although a dramatic increase in DNA quantity was observed with the change of deparaffinization agent, there were no noteworthy differences in the quality of DNA obtained. This may be due to major damage occurring during fixation protocols such as the extent of tissue autolysis, quality of fixative, duration of fixation, etc. which were not addressed in this study.

The DNA extracted from blood, fresh tissues, and FFPE tissues was bisulfite treated. FFPE tissues were initially treated with the Zymo EZ Methylation Gold kit but due to harsher bisulfite treatment and prolonged incubation cycle length (2.5 hours), they were also treated with the Zymo EZ lightning kit having a shorter conversion cycle length (60 mins) as treatment with Sodium bisulfite for a long duration largely impacted DNA quality. On estimating the DNA concentration post-bisulfite conversion fresh tissues had a DNA recovery of 80-85% post-conversion while degraded DNA from FFPE resulted in incomplete bisulfite conversion and lower DNA recovery even with the Zymo EZ lightning kit.

### 3.2 PCR Optimization and Sequencing

PCR conditions, including annealing temperature and primer concentration, were optimized for 11 epigenetic targets using gradient PCR. The resulting products were sequenced and high-quality data was obtained from fresh tissues, meeting recommended signal strength and quality score criteria (See Supplemental Figure 2). Compared to fresh tissues, FFPE tissues exhibited lower signal strength (See Supplemental Figure 3) due to high background noise, represented by a signal-to-noise ratio of less than 150, which is below the value of >150 recommended by Applied Biosystems (See Supplemental Table 2). To ensure accuracy, BigDye terminator kit control pGEM DNA was run alongside FFPE and fresh samples, detecting a total of 686 bases. Results showed good quality score values and sequencing quality, justifying the developed method (See Supplemental Figure 4). Therefore, it was found that the quality of DNA, along with the quantity, plays a major role in meeting sequencing data standards for accurate analysis of methylated nucleotides.

Given the low-quality sequencing data from the archived FFPE tissues resulting in the failed interpretation of methylation status at the target site, prospective cases of medulloblastoma were included in the study further. The developed methodology was applied and validated on surgically resected fresh tissues belonging to twelve cases.

Sequencing results covered 78% to 90% of amplicon length for most epigenetic targets, with exceptions for USP40, VPS37B, and RPTOR at 69%, 62%, and 66% respectively. Despite this, the target methylation site for all three targets was achieved, and optimum sequencing results were obtained for 10 out of 12 targets in the study. LHX6 was the only target that did not provide results. Nevertheless, each case had a successful interpretation of a minimum of 4/6 targets in classifier 1 and 3/5 targets in classifier 2 providing methylation status of 7 out of 11 targets per case.

### 3.3 Molecular subgrouping based on sequencing data

Chromas software (Technelysium) was used for trimming the low-quality reads from the sequencing results prior to their alignment with the reference sequence of each target using NCBI’s Global alignment tool. Figure 2 represents the chromatogram and sequence alignment for target cg02227036 in two cases clearly depicting the methylated (C to C) and unmethylated (C to T) status at the target methylation site. Each case was first tested for targets in Classifier 1 followed by Classifier 2 as per a study by Gomez et al. Based on findings in section 3.2, a criterion of successful data interpretation in 4/6 targets in Classifier 1 and 3 / 5 targets in Classifier 2 was decided to assign a molecular subgroup to cases. As a result, twelve prospective cases were successfully categorized into molecular subgroups, where results in two cases were found to be in accordance with the WNT group and thus they were not tested for Classifier 2, while five cases were subgrouped into Group 3 and five in Group 4 (see Table 1).

**Figure 2:**
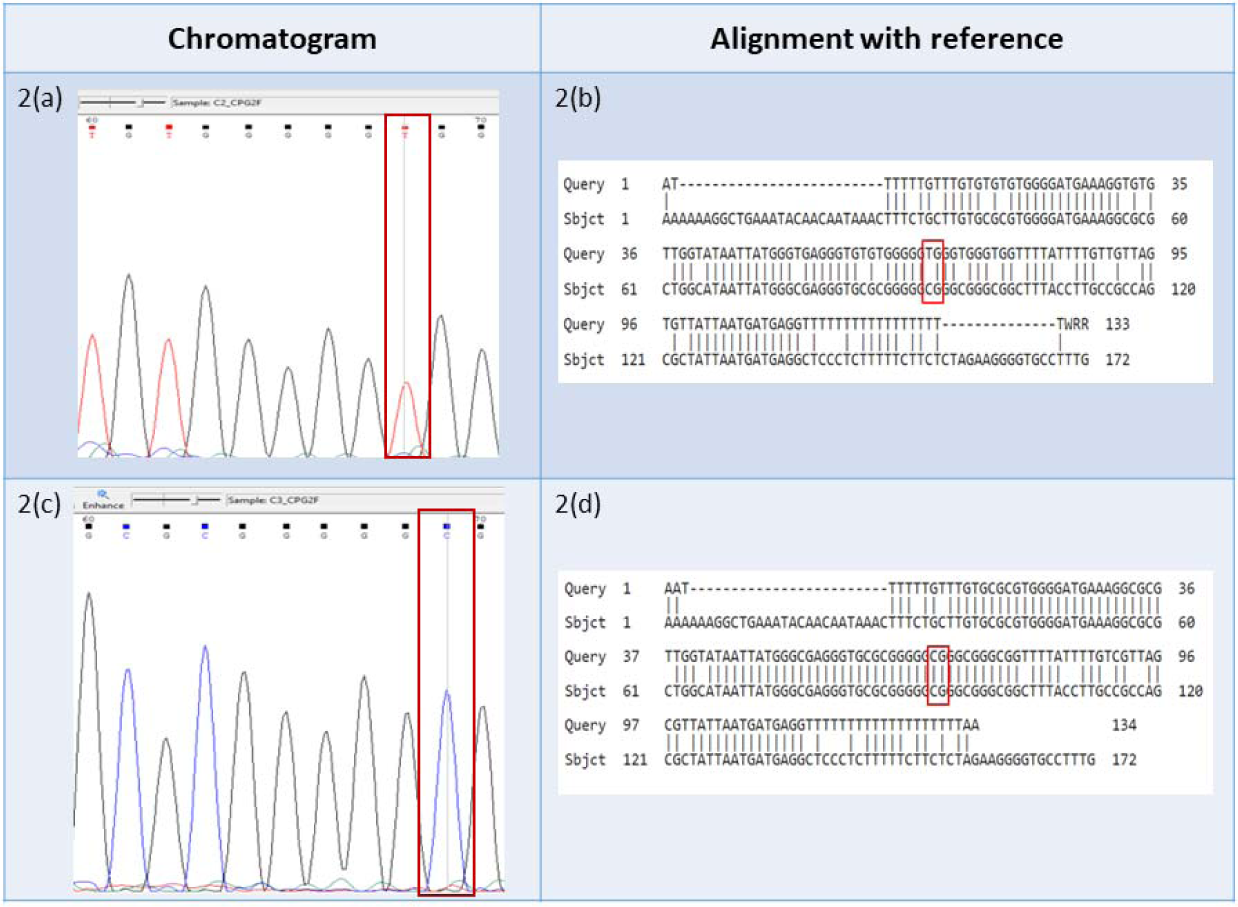
Figure representing chromatogram and sequence alignment for the target cg02227036 in two cases. Fig. 2(a) and 2(c) highlight the peak and signal obtained at the target site for two different cases. Fig. 2(b) and 2(d) depict the unmethylated (C to T) and methylated (C to C) status at the target methylation site in alignment with the reference.

**Table 1:**
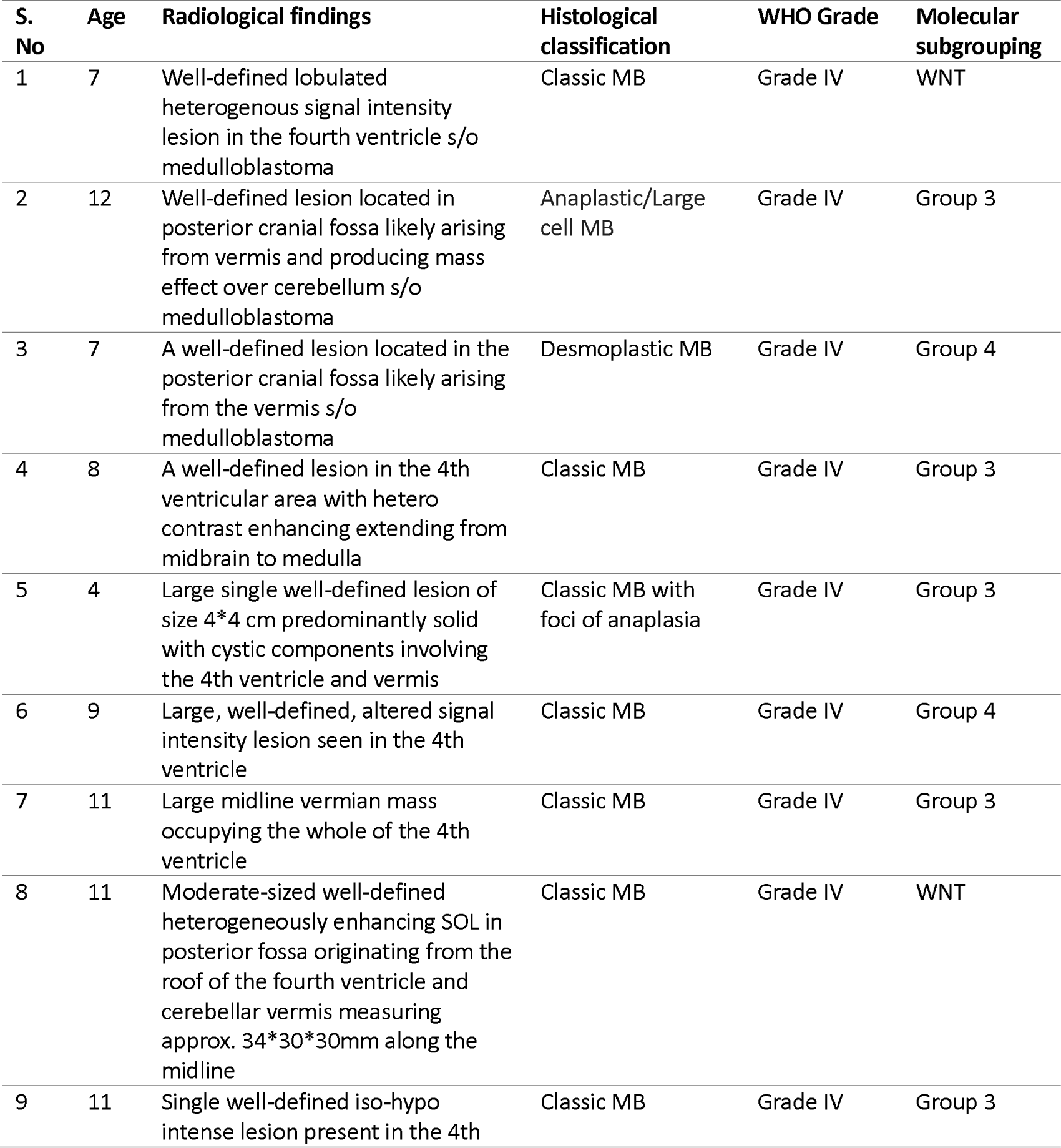
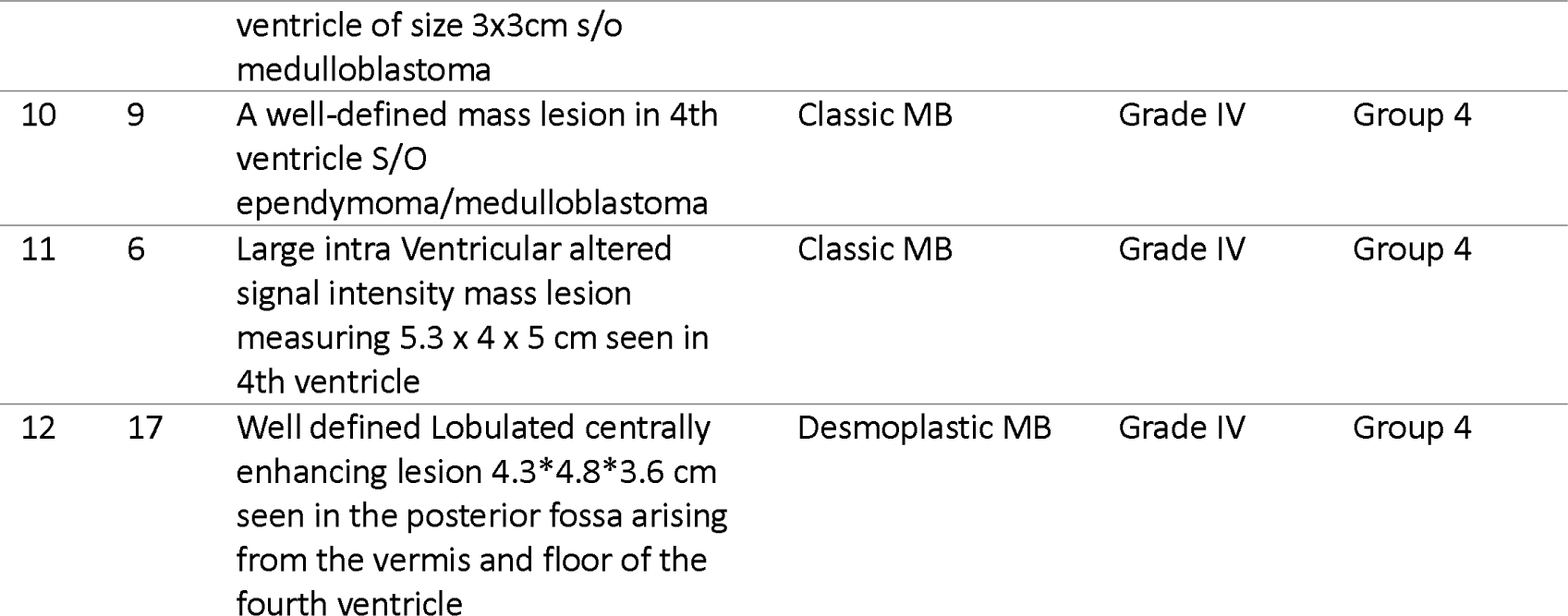
Table representing the clinical data of recruited cases and the molecular subgroups interpreted for the same using our methodology.

### 3.4 On correlating with Radiological and Histological features

Twelve patients with possible medulloblastoma identified by pre-operative radiology were included in the study. A confirmed diagnosis and histological classification were obtained post-operatively and correlated with molecular findings. The majority of cases (n=9) were classified as Classic histology. However, molecular subgrouping revealed 2 cases in the WNT group, 4 in Group 3, and 3 in Group 4. Two cases with desmoplastic histology were classified in Group 4 and one case with anaplastic histology was classified in Group 3 (see Table 1). Molecular subgrouping provided insight into prognosis and the need for clinical application.

## 4. Discussion

Medulloblastoma is the most common malignant CNS tumor in children and is associated with the most cancer-related deaths. In the past, its classification relied on histological and clinical findings, that were refined further by incorporating Immunohistochemistry however, the reproducibility of this method was not appreciable across many centers and also presented with a limitation of inability to classify Non-WNT/Non-SHH groups further into Group 3 and Group 4 (12). But advancements in molecular biology techniques now allow tumors to be classified molecularly into different groups. The updated fifth edition of the WHO classification of CNS tumors in 2021 recognizes and appreciates the inclusion of molecular diagnostics while retaining the histological approach to diagnosis and classification (7).

Despite the clear subgroups identified by genome and transcriptome profiling, their use in many clinical settings remains problematic. In contrast, methylation profiling in medulloblastoma tumors has proven to be a practical and highly effective method for classification following which numerous studies have reported methylation-based molecular classification (13–17).

One such study by Gomez et al. reported the development of a novel method for the classification of MB, based on identifying a novel set of epigenetic markers as classifiers (^Epi^WNT-SHH and ^Epi^G3-G4 classifier) by Linear discriminant analysis (LDA method) of DNA methylation microarray data from MB samples and their validation on frozen and FFPE samples by bisulfite pyrosequencing and direct bisulfite sequencing (18).

In this study, we used eleven epigenetic targets from previously mentioned classifiers to develop a fast and cost-effective method for classifying medulloblastoma into molecular groups based on methylation status. Sanger sequencing was utilized as it has been shown to be an effective method for methylation detection and quantification at a single nucleotide resolution (19–22). PCR-amplified epigenetic targets were sequenced to determine methylated/unmethylated status at the target methylation site. One target from Classifier 1 (LHX6) failed to provide sequences at the target region due to the close proximity of the target methylation site to the forward primer, resulting in failed interpretation. Poor-quality base calls often associated with the initial 50-100 bp sequences in Sanger sequencing could be the reason. Amplicon length also affects the read length of sequences as depicted by USP40 (167 bp; 69% coverage) and RPTOR (397 bp; 66% coverage) as shorter or longer fragments are often associated with mobility issues in capillary electrophoresis. However, similar issues were not reported by Gomez et al. in their study and thus the variation in results may be due to different experimental designs.

In terms of the feasibility of tissue type for sanger sequencing, we did not find satisfactory sequencing quality from FFPE samples procured from our institute. As supported by distinct literature, the fragmentation of nucleic acids induced by the duration of fixation and fixation step itself reduces the availability of desired amplifiable fragment length in FFPE DNA. Moreover, the amplification of PCR products greater than 200 bp is reported to be problematic in the case of FFPE samples (23,24). Several studies show age-based evidence of a negative impact on the quality and length of amplifiable DNA from long-stored tissue blocks (25–27). However, the tissue blocks we used were only 12-16 months old. Therefore, the difference in our result with other studies indicates that a greater degree of DNA degradation and fragmentation might have occurred at the fixation step, which differs from lab-to-lab protocol and processing not under the control of researchers. In addition, harsh chemical treatments in bisulfite conversion and necrosis associated with brain tumors may contribute to a certain degree of fragmentation. Accuracy-wise, sequencing of fresh tissue provided molecular data with optimum read length and quality score values compared to FFPE tissues, which may have altered molecular features by processing conditions, as reported in other studies (28). These studies suggested using a frozen tissue in addition to FFPE to check for concordance and accuracy (29).

The radiology reports, intra-operative impressions, and histopathological examinations provide only a diagnosis and do not address the prognosis. Our results show that we were able to obtain molecular subgroups with different prognoses for all recruited cases. Additionally, cases with similar histology were separated into distinct molecular subgroups. Therefore, adding molecular classification to routine radiology and histopathological examinations will aid in prognostication and improve patient care.

## 5. Conclusion

Diverse techniques exist for the molecular classification of medulloblastoma (MB). However, in developing countries, it is crucial to consider their feasibility and appropriateness. Here, we have developed and demonstrated the utility of PCR-based direct bisulfite sequencing for targeted methylation profiling of MB. This approach can be used in the majority of clinical settings and can provide clinicians with better prognostic information in addition to traditional clinical information obtained through radiology and histology.

## Acknowledgments

We would like to thank the Central Research Lab of SGPGIMS for their support and Dr. Saumya Sarkar for guiding the methodology of this study.

## Supplemental Data

**Supplemental Table 1:**
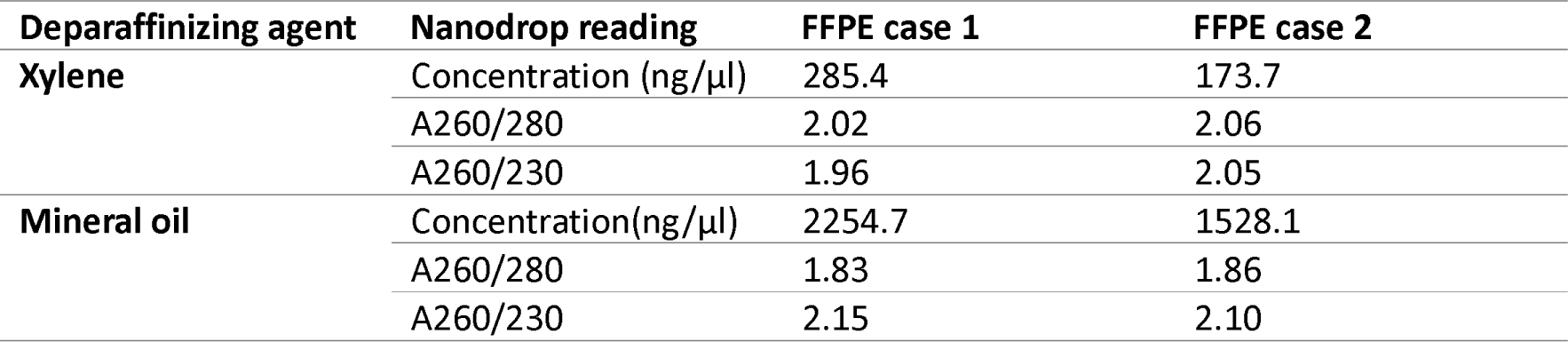
Nanodrop reading of DNA extracted from FFPE case 1 and FFPE case 2 depicting the effect of deparaffinization agent xylene and mineral oil on DNA yield.

**Supplemental Table 2.** shows the differences in signal intensity and signal-to-noise ratio for epigenetic marker CPG5F in two FFPE samples and a fresh sample. The fresh sample had higher signal intensity (>1000 RFU) and a higher signal-to-noise ratio (>150), as recommended by Applied Biosystems. The table also shows the number of bases detected and called by the KB base caller for all three cases.

| Samples | Average signal intensity |  |  |  | Signal-to-noise ratio |  |  |  | No. of bases detected |
| --- | --- | --- | --- | --- | --- | --- | --- | --- | --- |
|  | G | A | T | C | G | A | T | C |  |
| FFPE Case 1 | 560 | 931 | 1226 | 160 | 123 | 147 | 174 | 25 | 150 |
| FFPE Case 2 | 212 | 322 | 487 | 50 | 63 | 66 | 85 | 10 | 188 |
| Fresh tissue | 2098 | 1974 | 2528 | 177 | 523 | 327 | 371 | 30 | 188 |

**Supplemental Figure 1:**
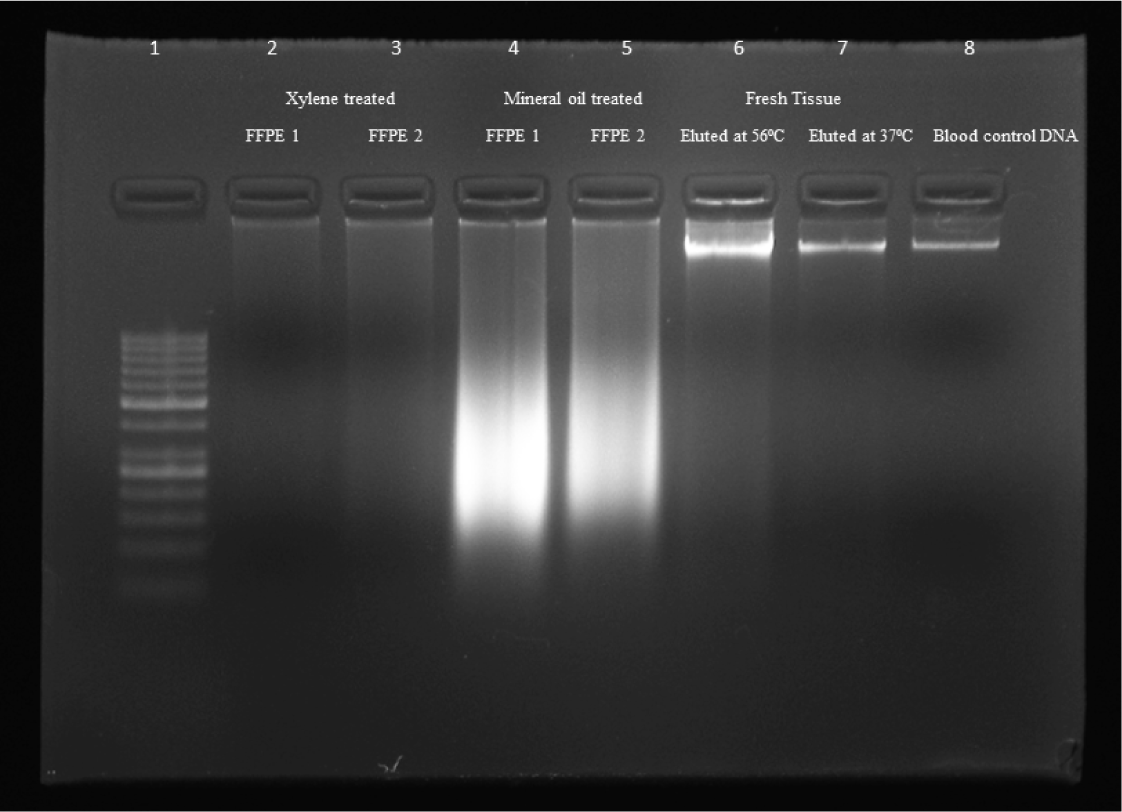
Gel doc image for DNA integrity analysis of DNA extracted from FFPE and fresh tissue. Lanes 2 and 3 show the smeared low-yield DNA extracted from two FFPE cases post-xylene treatment. Correspondingly, lanes 4 and 5 represent the high yield but less intact DNA post mineral oil treatment. Lanes 6 and 7 are represented by fresh tissue-derived DNA eluted at two different temperatures where elution at 56LJC shows mild smearing compared to elution at 37LJC. An intact band in lane 8 displays control blood DNA.

**Supplemental Figure 2:**
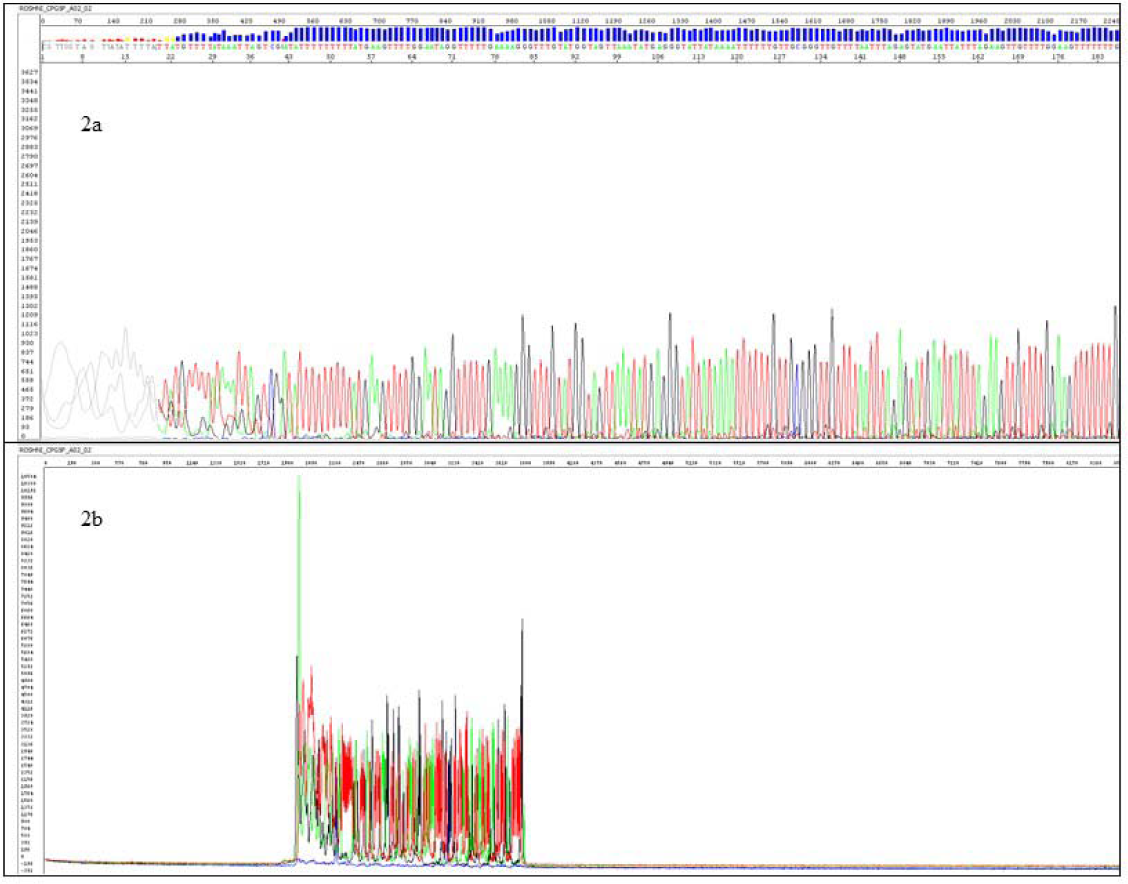
Electropherogram and raw signals of sequencing results from fresh tissue. Figure 2a depicts sequences, quality score values of each base called, and signal intensity in the case of fresh tissue. The blue bar on top of Fig. 2a represents the quality score value for each base identified. Figure 2b represents the overall raw signal intensity crossing the minimum threshold value of 1000 RFU.

**Supplemental Figure 3:**
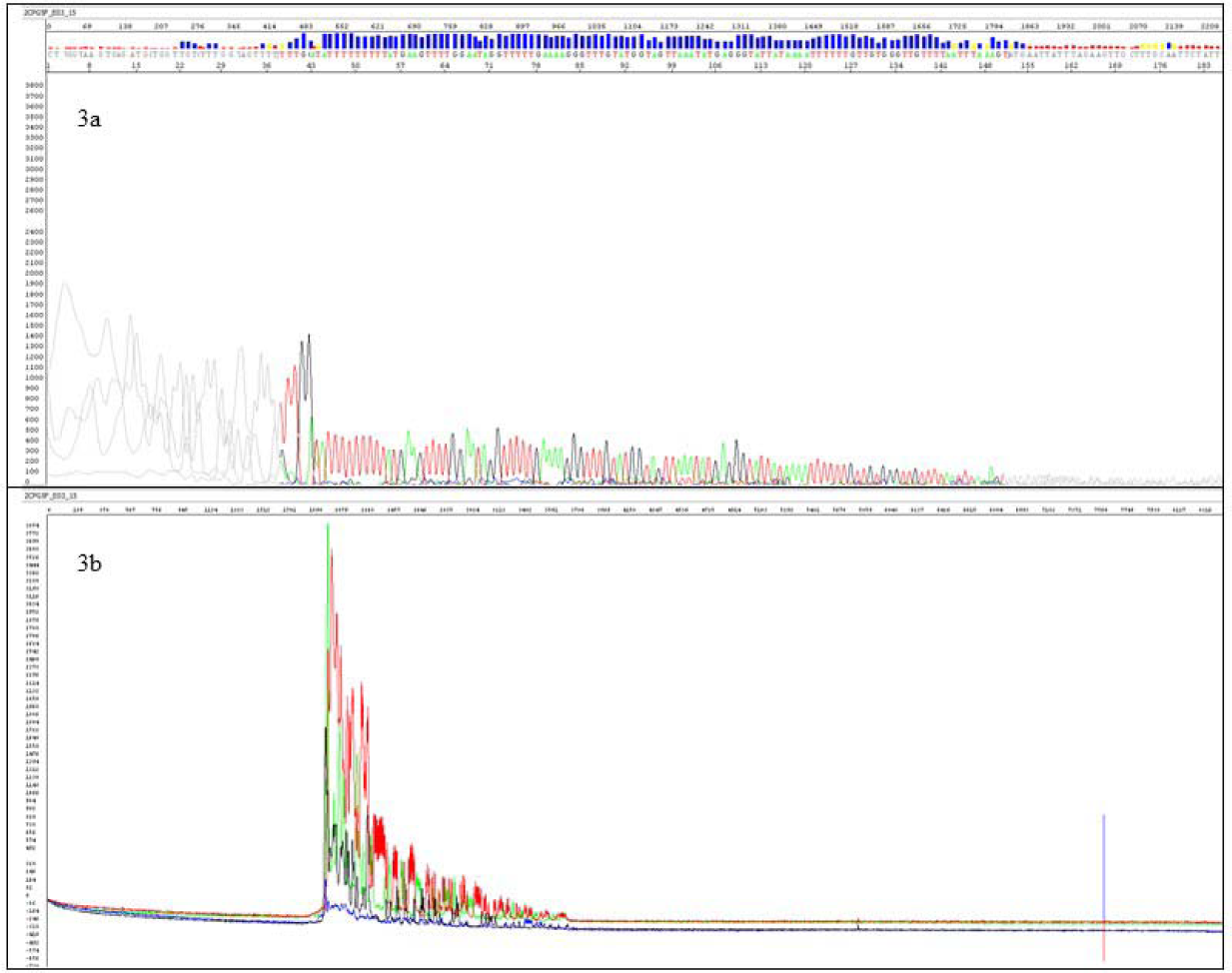
Electropherogram and raw signals of sequencing results from FFPE tissue. Electropherogram shows a poor-quality sequence due to low signal intensity and poor-quality score values in the case of sequencing from FFPE tissue (3a). Fig. 3b represents the raw signal intensity for the same

**Supplemental Figure 4:**
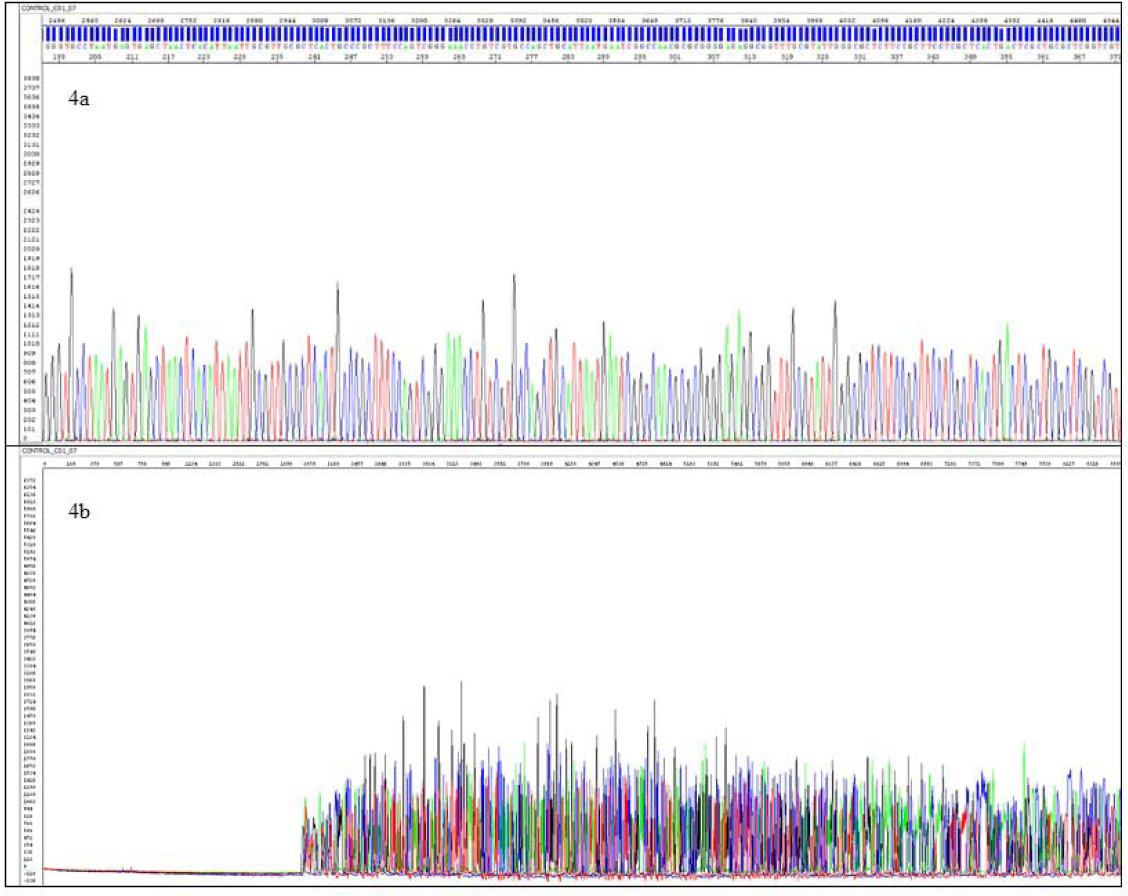
Electropherogram and raw signals of sequencing results from control. Electropherogram for pGEM control DNA using M13(–21) primer, representing high signal traces and quality score of bases (4a) and appropriate raw signal intensity (4b)

